# Behavior of downstream swimming brown trout in accelerating and high flow velocity - Movement matters

**DOI:** 10.64898/2026.04.22.720239

**Authors:** Falko Wagner, Ianina Kopecki, Jelger Elings, Ulf Enders, Andreas Lindig, Kathrin Maltzahn, Tom Roessger, Márcio S. Roth, Mansour Royan, Jürgen Stamm, Stefan Hoerner

## Abstract

Studies on active and sedated fish passing through turbines and pumps show different mortality and injury rates for both cases. Consequently, fish behavior appears to play a substantial role in these outcomes. However, direct behavioral observations in hydraulic machines using quantitative parameters to draw conclusions about the underlying mechanisms are hardly possible and remain understudied. In this study, we examined the behavior of adult brown trout (*Salmo trutta*) in an experimental flume under hydraulic conditions characterized by strong flow acceleration and high velocities typical of turbine and pump intakes. Fish movement behavior was analyzed based on a quantitative approach to enable the analysis of swimming behavior even in flow velocities exceeding the sprint swimming speed of fish. The application of Hidden Markov Models (HMM) to analyze activity states and movement modes of fish from video tracking data demonstrated significant effects of the spatial velocity gradient (SVG) and flow velocity on fish behavior. Notably, SVG emerged as the primary trigger for avoidance reactions when exceeding a threshold of 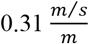. Fish exhibited distinct movement patterns under dark and daylight conditions, with more avoidance reactions in darkness. Whereas a considerable proportion of fish in daylight increased their swimming activity in the zone were flow velocity exceeded sprint swimming speed, in dark conditions no activity peak occurred in the same zone. The results illustrate how hydraulic conditions and lighting influence fish behavior. Integrating the behavioral rules identified in this study into numerical mortality-risk models could substantially improve their predictive accuracy. Thus, the findings allow for the development of less fish harming engineering solutions for hydropower facilities and pumping stations.

## 1. Introduction

Fish in heavily exploited rivers are exposed to a variety of man-made risks throughout their life cycle that can lead to injury or mortality. One of the significant threats comes from hydropower infrastructure, including turbines, pumps, and associated constructive elements. The injury and mortality risks during turbine passage have long been recognized and their quantification has been the subject of scientific study since the mid-20th century [1,2]. A meta-analysis by Radinger et al. [3] estimated the average mortality risk at hydropower facilities globally to be around 22%, though this varies widely between sites. Pumping stations and drainage systems, which are used to regulate water levels and manage flood risks, also pose considerable threats to fish [4,5]. These systems often entrain large numbers of fish with consequences analogous to those of turbines. Sea level rise due to climate change will lead to further expansion and modernization of pumping stations [6], increasing the need for effective fish protection measures. Regulatory authorities around the world are demanding fish protection strategies for both new and retrofitted hydropower plants and pumping stations [7–9].

The growing demand for fish-friendly technologies calls for innovative solutions that are both practical and effective. Currently, the evaluation of the injury and mortality risk of such technologies often involves experiments with live fish (e.g., [10–13]). By today these are considered the standard method [14,15]. However, alternative approaches for estimating fish mortality rates during turbine or pump passage emerge and are available. These include analytical [16], and numerical models [17,18], passive sensors [11,19], or combinations thereof [20,21]. While these methods cannot yet fully replace live fish tests as they can hardly quantify those risks in the required reliability, they offer valuable opportunities for comparing design alternatives of less fish harming turbines or pumps qualitatively and for estimating mortality risks as part of ethical considerations prior to conducting live fish experiments.

Nevertheless, fish behavior, an important factor influencing the risk of injury or mortality [22], is commonly not accounted for in alternative methods. In the uttermost cases it is simplistically assumed that fish drift passively with the flow through hydraulic structures. However, studies have shown that this assumption is not accurate, as active fish are more likely to be injured or killed than inactive individuals [23,24]. Initial approaches have been made to incorporate fish behavior into numerical models. A prerequisite for this, however, is a thorough understanding of how fish behave under flow conditions typical of turbines and pumps: These are in particular characterized by very high flow velocities which can become orders of magnitudes higher than the sprint capabilities of fish, strong pressure gradients, local flow accelerations and decelerations leading to significant spatial velocity gradients and high turbulent flows [25–27]. Quantitative values depend on hydraulic head and flow rates. Thus, they are site specific and vary largely.

The scientific community is still lacking fundamental knowledge about the behavioral responses of fish to hydrodynamic conditions [28], particularly during downstream movement in conditions where flow velocity exceeds the sprint swimming speed of fish. Downstream-migrating Atlantic (*Salmo salar*) smolts, exhibit strong rheotactic behavior often orienting to face the prevailing flow when encountering spatial velocity gradients (SVG) [29]. Such gradients are inevitable in turbines, pumps and their intakes. Chinook salmon smolts (*Oncorhynchus tshawytscha*) exhibited an avoidance response at the threshold of a mean SVG around 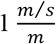 regardless of discharge [30]. Vowles and Kemp [31] have shown that brown trout (*Salmo trutta*) have a high probability of showing behavioral reactions if the SVG exceeds 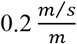 in the light and 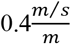 in the dark. Thus, the SVG appears to be an important parameter influencing swimming behavior of downstream migrating fish. However, the effect of flow velocities above sprint swimming speed on fish behavior as an isolated parameter has been poorly studied. In general, research on adult salmonid swimming behavior in these contexts is in its infancy [32].

Silva et al. [33] highlighted a critical gap in understanding fish passage, specifically how fish behavior and physiological responses are linked to local hydraulic conditions. The present study addresses this gap. An abstracted laboratory setup representative of the hydraulic conditions in pumps and turbine inlets, while not explicitly accounting for rotational flows within the machines, was developed. They are also valid for other human altered flows in hydraulic structures. The specific setup has been chosen to account for a broad variety of applications and to allow for an investigation of isolated parameters by examining the mutual behavioral effects of accelerated flow and velocities exceeding sprint swimming speeds in adult brown trout. The following two hypotheses have been formulated:

1. Downstream migrating fish do not behave passively when entering zones where the streamwise flow velocities exceed their maximum sprint swimming capacity. Fish continue to swim actively even when being subjected to involuntary drift.
2. Rapid acceleration of the streamwise flow velocity over short spatial distances (high spatial velocity gradients - SVG) induces significant changes in fish swimming behavior. The responses to hydraulic conditions can be modified by other stimuli such as illumination [31,34]. It can attract fish [35,36], but some studies also report mixed or contradictory effects [37]. Therefore, in our work we address the possible effect of lighting conditions by conducting experiments in both daylight and darkness leading to a third hypotheses:
3. Light condition affects the behavioral responses of fish to spatial velocity gradient and flow velocities which exceed the maximum sprint swimming capacity

Amaral et al. [38] provided first valuable insights into fish behavior in turbines from high-speed video observations. However, such studies of swimming behavior in high velocity environments are usually based on visual video observations without further tracking of motion or kinematics and are therefore qualitative. A strictly quantitative analysis of movement modes and swimming activity in the present study provides more detailed fish response data and valuable insights into fish behavior even when fish drift downstream due to high flow velocities exceeding the maximum sprint swimming speed of fish [39,40]. This may contribute to a more complete understanding of fish passage mechanisms and mortality risks.

## 2. Methods

### 2.1 Experimental Flume

To investigate the effects on fish behavior of flow velocity and SVG typical for conditions in turbine and pump intakes, a 30 m long and 2 m wide experimental flume was constructed. The maximum flow rate is 0.5 m^3^/s. The flume is composed of four sections, the filling tank (1), the partly open inlet section made of hard foam board for introducing the fish (2), a closed narrowing section with the observation area made of transparent acrylic glass (3) and an open outlet section (4) for recapturing the fish after the passage through the flume (Fig 1).

**Fig 1.**
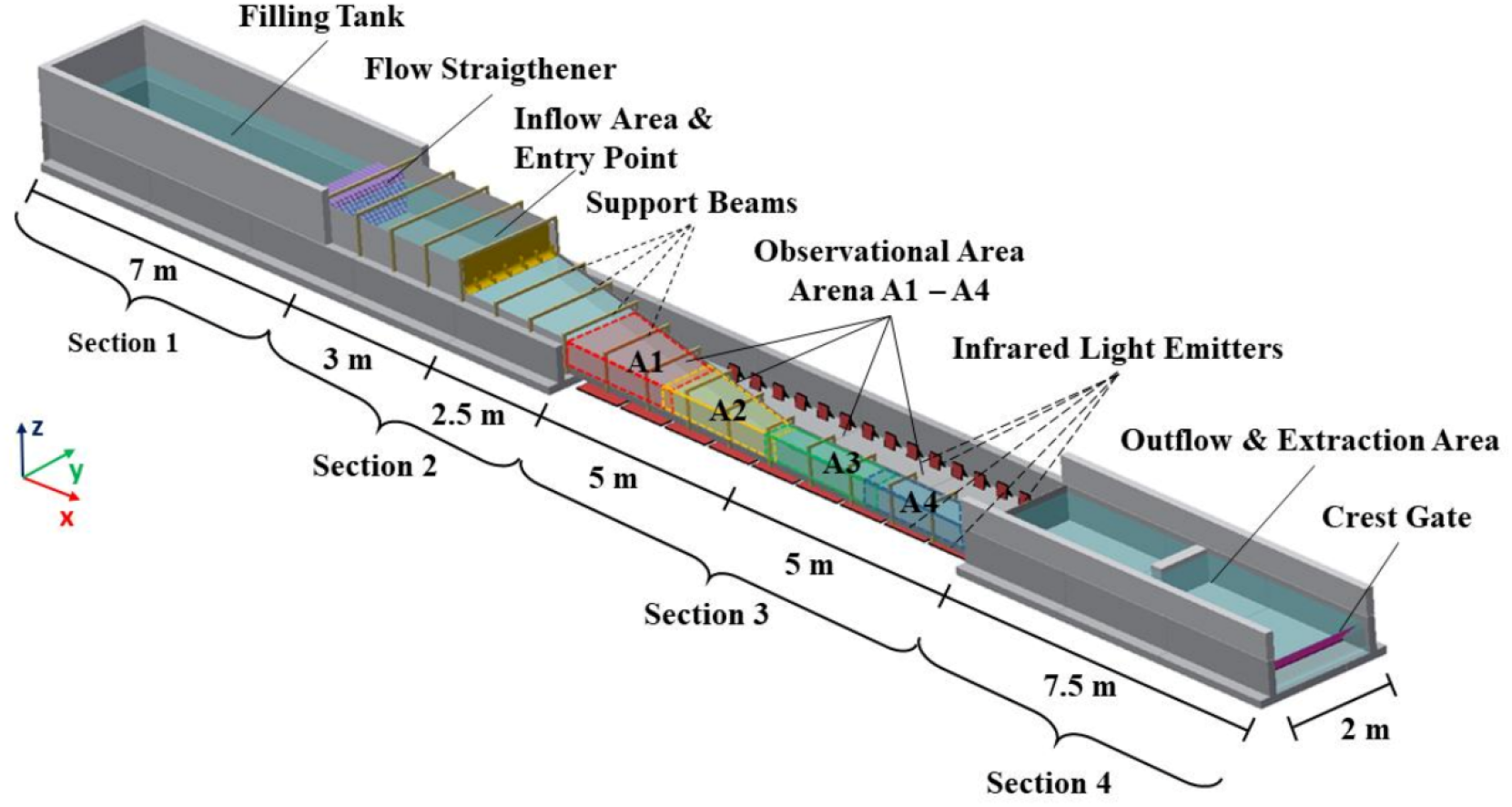
An isometric view of the experimental flume for live-fish experiments with its four functional sections (modified after [40]).

Within the observational area (Section 3), the flow cross-section of the experimental flume is reduced from 2 m to 0.45 m in width while maintaining a constant height of 0.4 m. This constriction results in an acceleration of the streamwise flow velocity (*v*_*x*_) up to approximately 3 m/s in arenas 3 and 4. The theoretical sprint swimming speed (*u*_*sprint*_) is exceeded for all experimental fish, calculated for a maximum 10 s swimming period based on fish length and water temperature for a rheophilic fish according to Ebel [41]. At the beginning of the high velocity flume part, the streamwise spatial velocity gradient (*SVG*_*x*_) decreases below the behavioral response threshold of 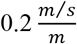 for brown trout [31]. This should allow the effect of flow velocity on swimming behavior to be studied independently from *SVG*_*x*_ in arena 3 and 4. Regarding the hydraulic properties, the observation area was designed to consist of four distinct zones, which differed in their hydraulic properties (Fig 2).

**Fig 2.**
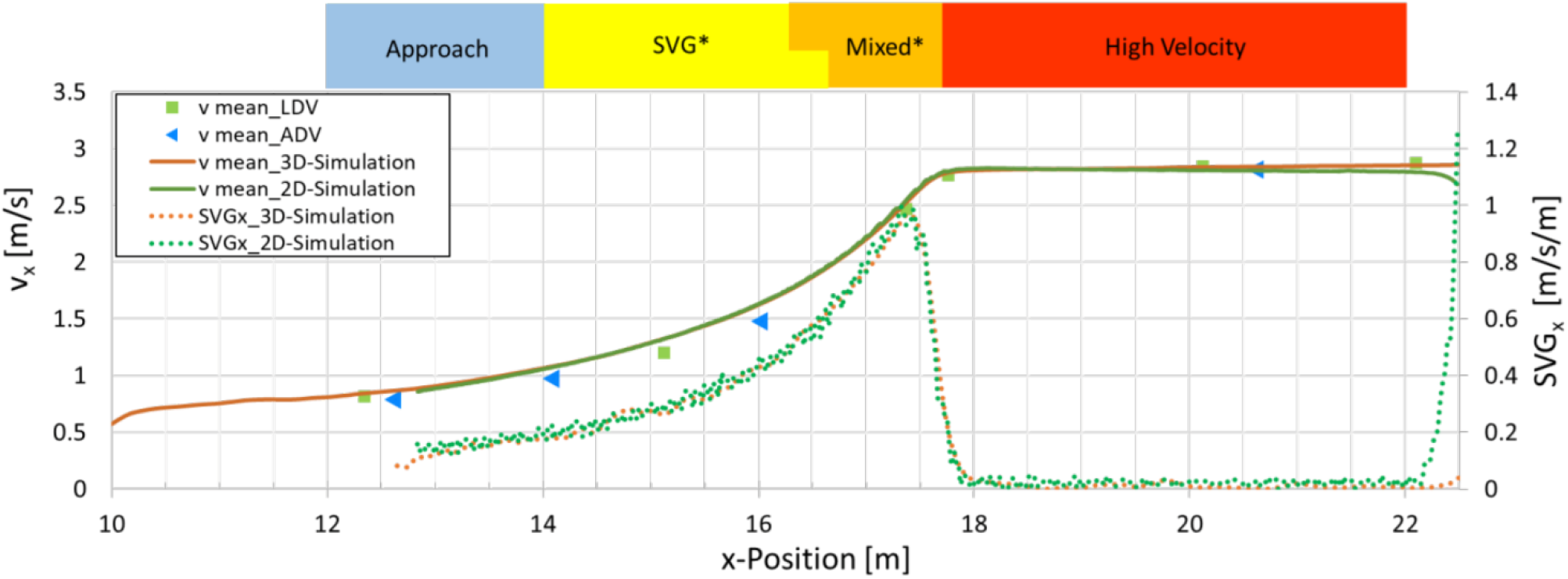
Simulated average velocity along the main flow vector (v_x_) (continues orange and green lines) and average SVG_x_ (dotted orange and green line) from 3D and 2D HN simulations in the observation section 3, measured average velocities v_x_ by ADV (blue triangles) and LDV (green squares), colored bars specify the four hydraulic zones (after [40]), Approach zone:-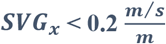; *v*_*x*_ < *u*_*sprint*_; *SVG*_*x*_ zone: 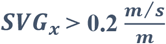; *v*_*x*_ < *u*_*sprint*_; Mixed zone: 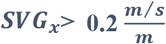; *v*_*x*_ ≥ *u*_*sprint*_; High velocity zone: 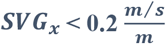; *v*_*x*_ ≥ *u*_*sprint*_; *gradual transition between the SVG and mixed zone due to the length dependent variation of the theoretical sprint swimming speed of each fish.

Approach zone: 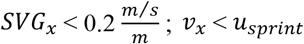

SVG zone: 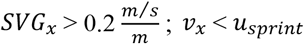

Mixed zone: 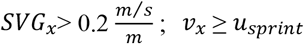

High velocity zone: 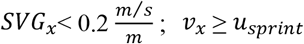

Variations in the theoretical sprint swimming speed of the fish (*u*_*sprint*_), depend on body length and water temperature. This causes a gradual transition between the SVG and mixed zone. The boundary between the SVG and the mixed zone was adjusted for subsequent behavioral analyses along the flume, based on the SVG threshold values identified in this study as indicating a high probability of avoidance responses by brown trout under either daylight or dark conditions. Consequently, the zoning used for the evaluation (Fig 7) did not completely correspond to that of the original flume design (Fig 2).

The comparison of simulations and flow velocity measurements (Fig 2) indicated that the flow in the observation area shows only minor vertical flow components and can be described by a steady two-dimensional (2D) hydrodynamic-numerical (HN) model with sufficient accuracy [36]. In consequence, 2D-HN simulations were performed with the HydroAS-2D model based on shallow-water equations [42]. The 2D modeled flow fields were used to calculate the relative fish speed by subtracting the modeled flow velocity from the absolute fish speed deduced from the tracking data [39]. The absolute local value of *SVG*_*x*_ component was calculated raster-based using the dominant flow velocity component *v*_*x*_ and a raster size Δ*x* of 2.5 cm:

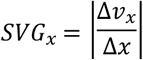

Δ*v*_*x*_= the difference in flow velocity between neighbor raster cells [m/s]

Enders et al. [30] and Vowles & Kemp [31] used following definition of *SVG* in their studies:

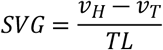

*v*_*H*_= flow velocity at the head of the fish [m/s]

*v*_*T*_= flow velocity at the tail of the fish [m/s]

*TL* = total length of the fish [m]

*SVG*_*x*_ represents a local equivalent of the *SVG* over total fish body used by [30] and [31] but eliminates a possible bias due to the fish body oriented not strictly along the main vector of the flow. Therefore, *SVG*_*x*_will be used in this study. Unless stated otherwise, SVG should be read as *SVG*_*x*_hereafter.

The 10*-*m*-*long observation area (Section 3) was monitored using eight infrared (IR) cameras (Basler AG, Germany; 1.3 MP) equipped with 4.4–11 mm lenses (Kowa Optimed Deutschland GmbH, Germany) and lens*-*mounted RG830 IR filters (Heliopan Lichtfilter*-*Technik Summer GmbH & Co. KG). Four cameras were mounted above the experimental flume and four on one side. The flume was fitted with actively illuminated IR backlight screens. Backlighting was provided by 40 IR spotlights and five custom*-*made IR*-*LED panels installed on the conduit bottom and on the left side facing downstream. The IR light was diffused using acrylic glass panels with diffuser sheets (300 μm thickness; light transmission 57% for the floor plate and 85% for side walls), producing a homogeneous and stable background illumination. This setup enabled reliable video tracking of fish as dark silhouettes against a bright background (Fig 3) under low-light conditions. [40]

**Fig 3.**
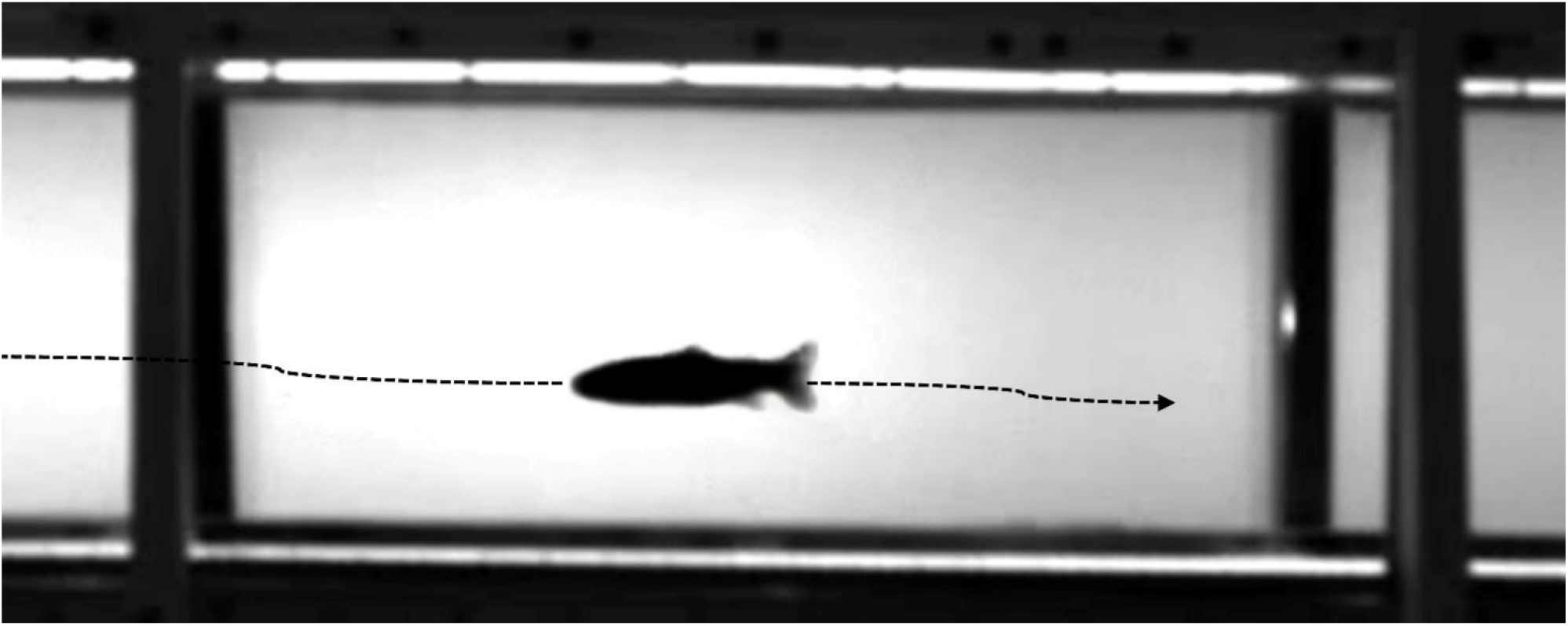
Image example from a lateral infrared camera; the fish is visible as a black silhouette against a light background. The dotted arrow visualizes the fish’s theoretical trajectory from the recorded video track.

The infrared light, with a wavelength of 850 nm, is not visible to brown trout [43,44]. According to Beach [45], infrared wavelengths greater than 750 nm can be used in fish experiments to simulate “dark” conditions without affecting behavior. Fish movements were recorded at a frame rate of 60 Hz, and trajectories were tracked using the software EthoVision XT (Noldus, Netherlands) [40]. This sampling rate corresponds to a spatial resolution with a tracking accuracy of < 5 cm, even under conditions of flow velocities approaching 3 m/s.

### 2.2 Fish handling

A total of 98 wild brown trout were captured by electrofishing in the river Apfelstädt in March 2021. Prior to the experiments, the fish were held for at least 48 hours under natural light conditions in six 500 L tanks at the experimental facility (maximum stocking density: 8.6 kg/m^3^). Throughout the holding period and during all flume experiments, activities that could induce vibrations, shocks, loud noises, or abrupt changes in lighting within the experimental hall were strictly avoided. Water in the holding tanks was aerated and continuously recirculated through a mechanical filter combined with UV and ozone treatment. Daily replacement of at least 10 % of the total tank volume ensured consistently high water quality and stable temperature conditions. Both the holding tanks and the experimental flume were supplied with water from the facility’s deep storage tank, ensuring identical water temperature and chemical parameters across systems. Dissolved oxygen concentration, pH, conductivity, nitrate, nitrite, chloride concentration and temperature were checked twice a day to monitor the water quality. All fish spent a maximum of 14 days in the experimental facility before being released into their original water bodies.

### 2.3 Fish Experiments

A subset of 54 individuals with a uniform mean body length of 0.262 m (Wilcoxon rank-sum test: W = 462, p = 0.0911; Shapiro–Wilk test for normality: p < 0.005) was selected for behavior analysis, comprising 28 fish from the daylight trials and 26 from the dark trials (Table 1). This size-matched subgroup was a requirement to exclude body length as a potential confounding factor influencing swimming behavior.

**Table 1.** Lighting conditions, number of fish, total length (TL), fish mass, it standard deviation (SD), water temperature and theoretical sprint swimming speed (maximum swimming duration of 10 s of a rheophilic fish [41] of the experimental fish in the flume experiments.

| **Cond.** | **Mean**<br>**Light**<br>**[Lux]** | **N**<br>**fish** | **Mean**<br>**TL**<br>**[m]** | **SD of**<br>**TL [m]** | **Mean**<br>**fish**<br>**mass [g]** | **Mean**<br>**WT**<br>**[°C]** | **Min.**<br>**WT**<br>**[°C]** | **Max.**<br>**WT**<br>**[°C]** | **Mean**<br>**sprint**<br>**speed**<br>**[m/s]** | **Min.**<br>**sprint**<br>**speed**<br>**[m/s]** | **Max.**<br>**sprint**<br>**speed**<br>**[m/s]** |
| --- | --- | --- | --- | --- | --- | --- | --- | --- | --- | --- | --- |
| **Light** | 84.5 | 28 | 0.261 | 0.017 | 156 | 10.4 | 10.0 | 11.0 | 1.88 | 1.78 | 2.21 |
| **Dark** | 36.7 | 26 | 0.263 | 0.013 | 160 | 10.7 | 10.0 | 12.0 | 1.94 | 1.67 | 2.04 |
| **Total** | 59.8 | 54 | 0.262 | 0.015 | 158 | 10.5 | 10.0 | 12.0 | 1.91 | 1.67 | 2.21 |

A subset of 54 individuals with a mean body length of 0.262 m—showing no significant difference between groups in daylight and dark experiments (Wilcoxon rank-sum test: W = 462, p = 0.0911; Shapiro–Wilk test: p < 0.005) was selected for the behavior analysis. The size-matched subgroups consisted of 28 fish in the daylight trials and 26 in the dark trials (Table 1). Controlling for body length was necessary to exclude size as a potential confounding factor affecting swimming behavior.

The flume experiments were conducted between 8th and 24th of March 2021 during the natural downstream migration season of local brown trout. To create dark conditions, an enclosure made of an aluminum frame covered with blackout fabric was installed around the entire experimental setup, including the holding tank and handling area [40]. This reduced mean illumination to 36.7 lux, compared with 84.5 lux under daylight conditions (Table 1). Water temperature during the trials ranged from 10 °C to 12 °C. The theoretical mean sprint swimming speed of the brown trout during the experiments was 1.91 m/s (Table 1). Detailed information on fish length, mass, water temperature, and corresponding sprint speeds under light and dark conditions is provided in Table 1.

During the experiments, one single adult brown trout was tested per trial. Each fish was individually transferred in a closed 15 L bucket from its holding tank to the entry point in section 2 of the flume (Fig. 1). The flow velocity at the entry point was kept below 0.4 m/s, enabling the fish to enter the observation section voluntarily. Each trial continued either until the fish traversed the entire flume or until a maximum duration of 45 minutes was reached. After each trial, the fish was gently netted and returned to its holding tank. To avoid potential learning or habituation effects, each individual was used only once.

All procedures were conducted in accordance with animal welfare regulations and approved by the responsible authority of the federal state of Saxony, Landesdirektion Sachsen (permit number TVV 2/2020), in compliance with European animal welfare legislation (permit number TVV 2/2020) according to the European laws on animal welfare [46].

### 2.4 Assessment of fish behavior

Under high-velocity flow conditions movements and transitions can occur too rapidly to be reliably detected by visual observation. An accurate characterization of fish behavior requires the use of quantitative, algorithm-based metrics. To address this challenge, a high-resolution analytical framework was applied, enabling behavioral assessment at spatial and temporal scales suitable for the dynamic hydraulic environment of the flume [39,40,47].

The behavioral analysis consisted of three core components: activity, movement, and reaction (Fig. 4). Each component was derived from automated tracking data and quantified using mathematical algorithms, ensuring objective, reproducible, and observer-independent evaluation [40]. This approach allows for consistent interpretation of behavioral patterns and facilitates direct comparison across experimental conditions, setups, and even fish species, in particular when behavioral adjustments are subtle or not visually discernible.

**Fig 4.**
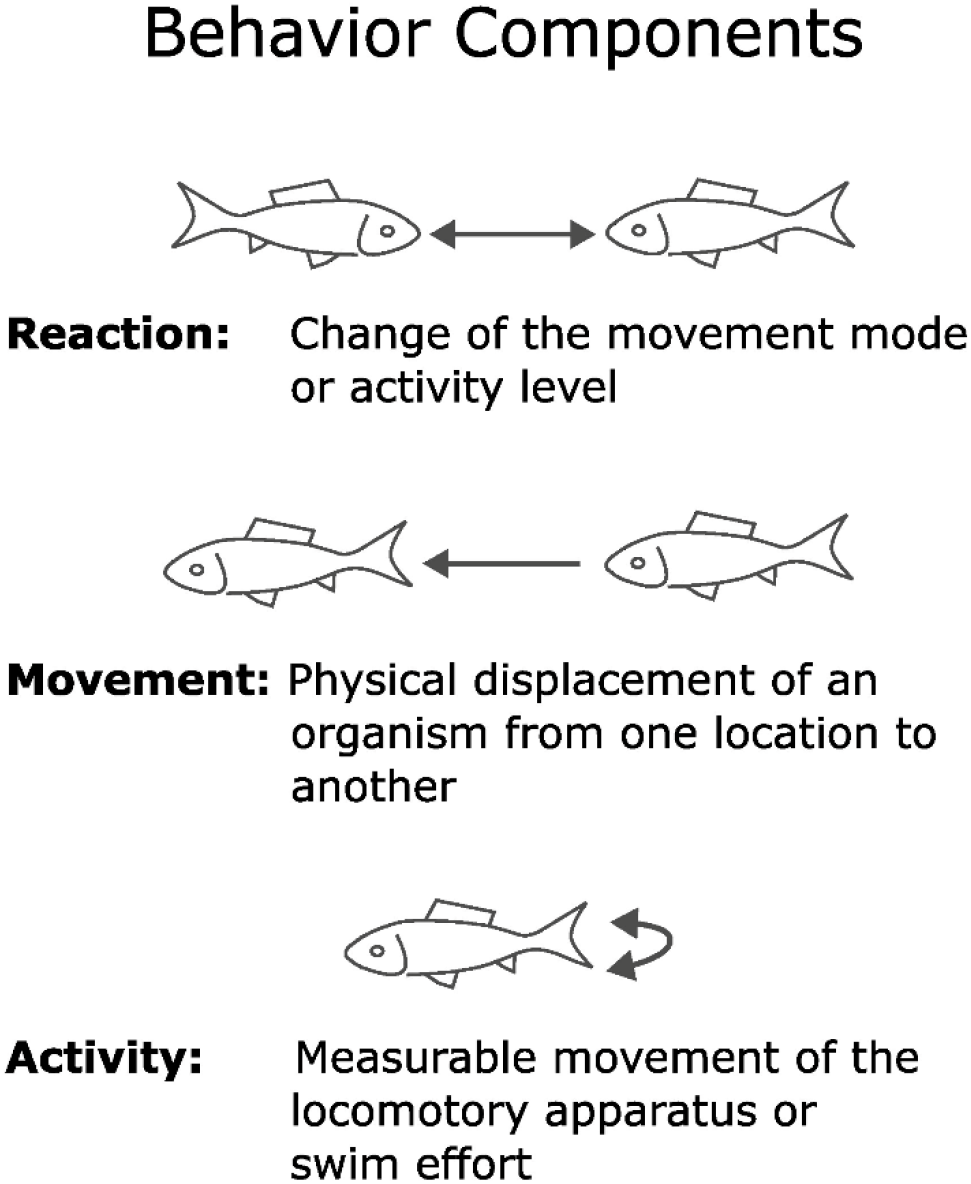
Components used for quantitative characterization of the fish behavior in the ethohydraulic experiments and the component definitions.

To distinguish between different movement modes, the relative swimming speed (*u*_*rel*_) and its ratio to the flow velocity (*v*_*x*_) were used as key parameters (Fig. 4). These metrics enabled the classification of five basic movement modes, each describing the fish’s displacement direction and rheotactic orientation during a given time step.

Although fish activity can be visually estimated by observing locomotor effort, such as tail-beat intensity in body– caudal-fin swimmers an objective and reproducible assessment demands quantitative metrics. Parameters such as tail beat frequency, swimming speed, and swimming power provide robust quantitative indicators of swimming activity.

In this study, fish activity was quantified from tracking data by calculating normalized swimming power (*P*_*norm*_, Formular 1), following the approach of Kopecki et al. [39]. *P*_*norm*_ expresses the fish’s swimming effort relative to its theoretical sprint swimming power (*P*_*sprint*_), which is calculated based on body surface area [48] the guild-specific theoretical sprint swimming speed (*u*_*sprint*_) influenced by water temperature [41], and a drag coefficient of 0.008 as defined by Kopecki et al. [39]. Swimming performance was conceptually divided into two components

> Power to overcome flow resistance (*P*_*drag*_),
>
> Power required for acceleration (*P*_*acceleration*_)

However, previous analyses confirmed that *P*_*acceleration*_ was negligible in all cases compared to *P*_*drag*_, allowing the focus to remain on the drag component as the dominant factor in swimming power estimation [37].

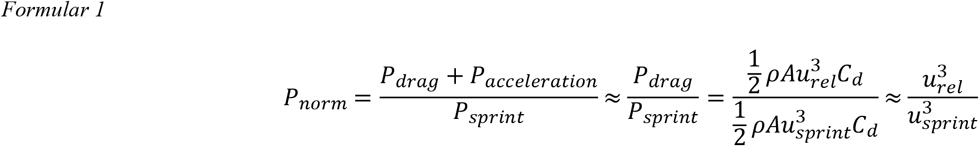

ρ is the density of water [kg/m^3^],

*A* is the fish surface area [m^2^], *A* = 0.401*TL*^2^ see [48]

*C*_*d*_ is the drag coefficient, assumed constant for lower and sprint speed, see [39] [-]

In arenas 3 and 4 of the experimental flume, flow velocities exceeded 2.5 m/s (Fig. 2), surpassing the theoretical maximum sprint swimming speed of brown trout (2.21 m/s; Table 1). Under such conditions, fish are theoretically unable to hold their position or escape upstream, making upstream avoidance responses undetectable within the five predefined movement modes (Table 2). Nevertheless, within the movement mode drift controlled, where fish are advected downstream by the flow, a marked increase in swimming effort can still occur. Such elevated effort represents an active behavioral response aimed at resisting, moderating, or delaying involuntary downstream drift despite the overwhelming hydraulic forces.

**Table 2.**
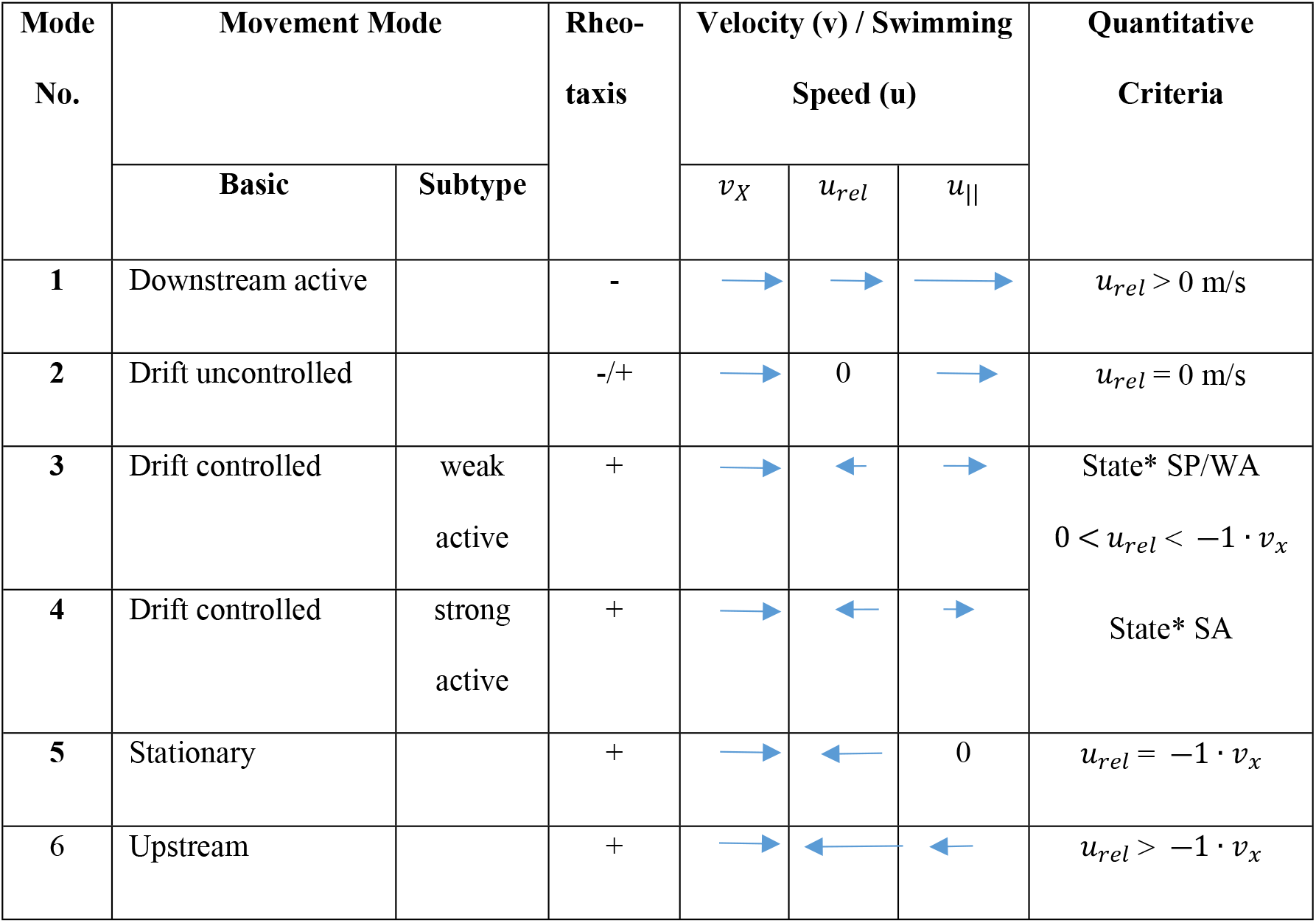
Movement modes of fish in the ethohydraulic tests as a function of the relative swimming speed (u_rel_), the flow speed along the x vector (v_x_) resulting in the absolute swimming speed (u_||_) and the two subtypes for drift controlled based on the activity states from [52], with weak swimming activity consisting of the *states by the HMM model defined states SP = semi passive, WA = weak active and SA = strong active; SP and WA both representing weak swimming activity without substantial delay of downstream drift, SA represent strong activity delaying the downstream movement.

To detect these subtle changes in swimming activity, Hidden Markov Models (HMMs) were applied using normalized swimming power (*P*_*norm*_) as the primary input variable. HMMs infer unobserved (latent) behavioral states from observed movement data by modeling state transitions over time. In this study, they were used to classify fish behavior into discrete activity states, distinguish between low and high swimming effort within the drift controlled mode, link these states to their spatial distribution, and examine transitions in relation to the flow parameters (flow velocity and SVG) experienced by the fish.

Model fitting was conducted using the *momentuHMM* package [49]. Based on initial data inspection, three behavioral states were considered: semi-passive (SP), weak activity (WA), and strong activity (SA). To reduce the risk of local maxima, each model was fitted 50 times with randomly assigned initial values [50]. The model with the highest log-likelihood was selected for further analysis. Behavioral states were then assigned to each fish position using the Viterbi algorithm [51], enabling the identification of meaningful shifts in swimming effort that would otherwise be masked under the assumption of passive drift. Both SP and WA represent low-power swimming not effectively slowing downstream movement within the drift controlled mode. In contrast, SA reflects substantially increased swimming effort that effectively reduces drift speed.

Accordingly, the drift controlled movement mode was subdivided into two behavioral subtypes:

- Weak activity: comprising SP and WA
- Strong activity: corresponding to SA

This refinement led to an expanded behavioral classification scheme comprising a total of six movement modes (**Table 2**), allowing for a more detailed interpretation of fish responses under high velocity conditions.

A transition from one of the six defined movement modes to another can be interpreted as a behavioral reaction of the fish to either intrinsic or extrinsic stimuli and is referred to as a reaction (Fig 4). According to the classification scheme outlined in Table 2, a switch from a movement mode with a lower to a higher mode number generally corresponds to an increased delay in downstream movement. Such transitions are therefore considered indicative of an avoidance response, reflecting the fish’s attempt to resist or mitigate downstream movement caused by the flow.

### 2.5 Statistical analysis

Generalized Additive Models (GAMs; [48]) were used to assess the influence of hydraulic and environmental predictors on the avoidance reaction rate along the x-axis of the flume. Predictor variables included the spatial velocity gradient in the x-direction (*SVG*_*x*_), the streamwise flow velocity (v_x_), and lighting condition. The avoidance reaction rate was calculated for each of the 38 virtual flume segments along the x-axis (25 cm width) as the number of behavioral reaction events (i.e., transitions to movement modes with higher numerical values) relative to the total number of fish observations in that segment.

To account for potential nonlinear effects and interactions among predictors, the GAMs included smooth main effects (s) and their respective two-way tensor product interactions (ti). Furthermore, to address the potential over-parameterization given our specific sample size (n=38), we applied thin plate regression splines with an added shrinkage penalty (bs = “ts”) [53].

> The full model was specified as:
>
> avoidance_reaction_rate ∼
>
> s(SVGx, bs = “ts”) +
>
> s(v_x, bs = “ts”) +
>
> s(lighting condition, bs = “ts”) +
>
> ti(v_x, SVGx, bs = “ts”) +
>
> ti(v_x, lighting condition, bs = “ts”) +
>
> ti(SVGx, lighting condition, bs = “ts”)

To refine the model structure, we implemented an iterative selection procedure to optimize the basis dimension (k) for each smooth term. We initially set *k* = 10 to provide sufficient flexibility for capturing potential nonlinear relationships. However, this value increased the risk of overfitting. Therefore, we evaluated model diagnostics using the gam.check() function in the mgcv package (version 1.9-1; [54]) in R [55], along with the effective degrees of freedom (EDF) and associated *p*-values. Based on these assessments, the basis dimension for all smooth terms was reduced to *k* = 5. This choice maintains the ability to detect nonlinear patterns while preserving degrees of freedom, which is particularly important given the relatively small sample size (*n* = 38) [55].

This formulation allowed flexible estimation of both main effects and interactions while preventing over smoothing and ensuring identifiability of component smooths. The same model but without the variable lighting condition and its interactions with hydraulic variables were applied for the dark and daylight datasets separately.

To ensure robust estimation of model parameters and smooth terms, all Generalized Additive Models (GAMs) in this study were fitted using the Restricted Maximum Likelihood (REML) method. REML is a reliable approach for estimating smoothing parameters in GAMs, particularly in complex models involving multiple smooth terms and interactions [56]. Unlike traditional maximum likelihood estimation, which simultaneously estimates fixed effects and variance components, REML focuses specifically on the estimation of variance components such as smoothing parameters by maximizing the likelihood of a transformed dataset that excludes fixed effects. This separation reduces bias in the estimation process and enhances model stability. In this study, REML was chosen for three primary reasons:

1. Optimizing smoothness of fitted curves for key predictors such as (*SVG*_*x*_) and v_x_ under both illumination conditions.
2. Preventing overfitting by penalizing unnecessarily complex smooth terms that do not substantially improve model fit.
3. Improving generalizability, ensuring stable model performance across flume segments and individual fish.

The REML-based fitting was implemented using the mgcv package (version 1.9-1; [54]) in R [55], which automatically selects appropriate smoothing parameters via penalized likelihood. This approach provided a reliable and interpretable framework for modeling the relationships between hydraulic and biological predictors and fish behavioral responses. For variable selection, the models incorporated a shrinkage-based penalty that allows individual smooth terms to be reduced effectively to zero when unsupported by the data, thereby removing them from the model structure [53]. All models used a Beta Regression with logit link function. A base model was constructed by smoothing the response variable over the x-coordinate of the flume. In daylight dataset the response variable contains zero values which the standard Beta distribution will not support. To accommodate the logit link function in the GAM, a small constant (epsilon) of 0.0001 was added to the response variable for all observations. This transformation ensures all values are strictly between 0 and 1 while preserving the relative distribution of the data and is a common approach for handling boundary values in Beta regression [57].

To identify thresholds for *SVG*_*x*_ and *v*_*x*_ that predict high reactions probabilities, a recursive partitioning approach was applied to compute conditional inference trees [58]. These were computed with the ctree() function from the R package ‘partykit’ (version 1.2.22; [59]) in R (v. 4.4.1; [60]). Conditional inference trees use statistical significance testing to iteratively partition the dataset into mutually exclusive and internally homogeneous groups, based on the most informative explanatory variable at each step. This method avoids overfitting and selection bias by separating variable selection from split determination, thereby identifying predictors that best explain variability in the response variable [58].

To test for differences in avoidance reaction rates between fish in daylight and dark experiments, a Mann–Whitney U test was conducted. This nonparametric test was chosen because the data did not meet the assumptions of normality, as confirmed by the Shapiro–Wilk test (p < 0.05).

## 3. Results

The behavioral analysis using Generalized Additive Models (GAMs) revealed significant effects of hydraulic variables on the avoidance reaction rate, defined as transitions to movement modes associated with increased delay in downstream movement. The probability of avoidance reactions was significantly influenced by the spatial velocity gradient (*SVG*_*x*_) (edf = 1.0019, F = 32.82, p < 0.0001) and the streamwise flow velocity (v_x_) (edf = 0.8308, F = 4.904, p = 0.0116). No significant effect of lighting condition (daylight vs. dark) was detected in the pooled dataset.

When analyzing daylight and dark conditions separately, using a reduced model with *SVG*_*x*_, v_x_, and their interaction as predictors, both variables remained significant:

Under dark conditions, both SVG_x_ and v_x_ were significant as individual predictors:

> *SVG*_*x*_: edf = 0.7847, F = 3.875, p = 0.0207
>
> v_x_: edf = 0.8109, F = 4.159, p = 0.0204
>
> v_x_ × SVG_x_ interaction: edf = 0.3788, F = 0.641, p = 0.1722

Under daylight conditions, SVG_x_ was significant as a single predictor, while v_x_ showed significance only in interaction with SVG_x_:

> *SVG*_*x*_: edf = 0.8134, F = 3.938, p = 0.01320
>
> v_x_ × SVG_x_ interaction: edf = 1.0197, F = 5.635, p = 0.0068
>
> v_x_: edf = 0.0.0001, F = 0.000, p = 0.5388

These results suggest that *SVG*_*x*_ consistently plays a dominant role in triggering avoidance behavior, while the influence of flow velocity varies with lighting conditions. In daylight its effect depends on the magnitude of the spatial velocity gradient.

The strong influence of *SVG*_*x*_ on fish swimming behavior was further confirmed through conditional inference tree analysis, which examined avoidance reaction rates in the 38 flume segments along the x-axis. Among the predictor variables flow velocity, SVG_x_, and lighting condition the *SVG*_*x*_ consistently emerged as the most decisive factor.

For the pooled dataset (combining daylight and dark experiments), *SVG*_*x*_ formed the first and most significant node, resulting in a clear separation between groups with low and high avoidance reaction rates. A similar pattern was observed when analyzing the daylight data independently. In both cases, the optimal threshold for group separation was identified at an *SVG*_*x*_ value 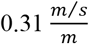 (Fig 5 and Fig 6).

**Fig 5.**
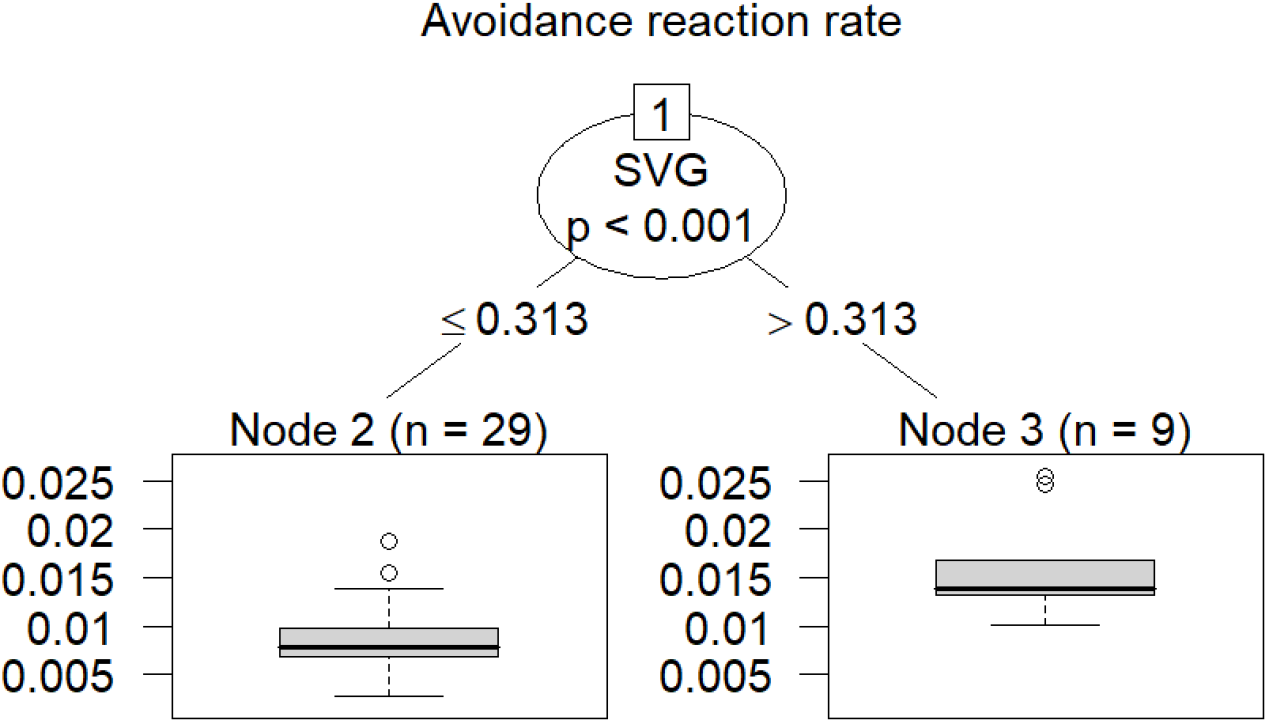
Boosted regression tree for predicting the reaction rate of avoidance behavior (delay of downstream movement) of brown trout in the experimental flume based on the independent variables flow velocity (*v*_*x*_), spatial velocity gradient (*SVG*_*x*_), lighting conditions and fish length.

**Fig 6.**
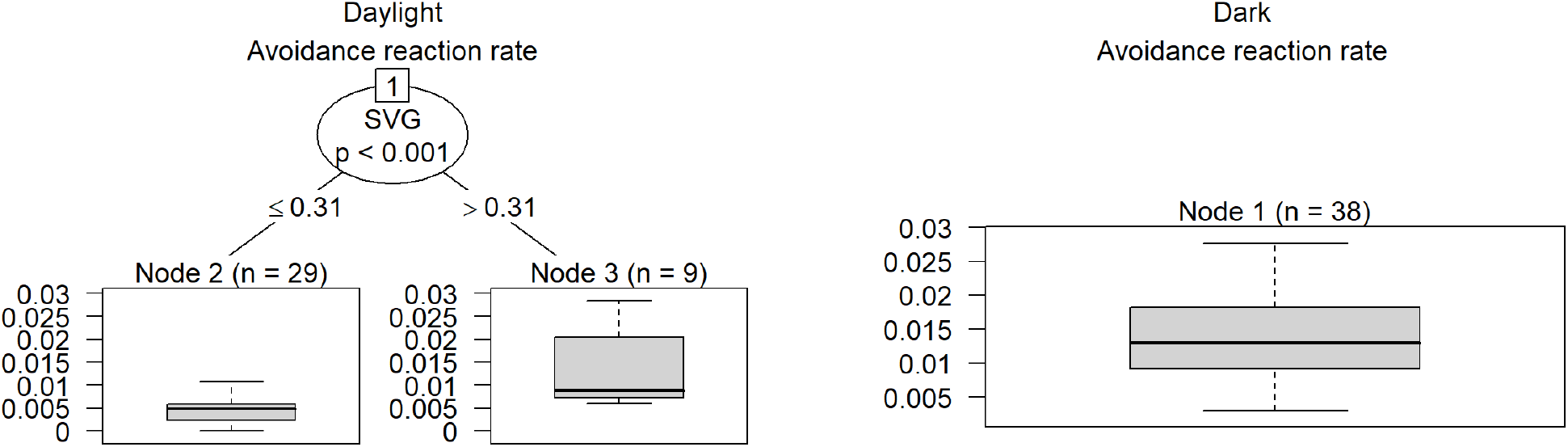
Boosted regression tree predicting the reaction rate within 25 cm wide virtual flume volume segments along the flume x-axis for avoidance behavior (delay of downstream movement) of brown trout in the experimental flume based on the independent variables (*v*_*x*_), spatial velocity gradient *SVG*_*x*_and fish length, in daylight (left) and dark (right) experiments.

In contrast, the analysis of the dark condition data did not yield a statistically significant split or threshold for any of the predictor variables. This suggests that under low-light conditions, fish avoidance behavior may be influenced by additional more complex factors, making threshold identification less conclusive (Fig 6).

The avoidance reaction rate within the 38 virtual flume segments varied along the x-axis of the experimental flume under both lighting conditions (Fig. 7). However, the variation was greater under daylight than in darkness Fig. 7). In the daylight experiments, the strongest increase of avoidance reaction rate began at position x ≈ 15 m (Fig. 7), where the specific gradient of velocity (*SVG*_*x*_) exceeded 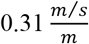, and the flow velocities being well below the fish’s maximum sprint swimming speed (Fig. 2). The reaction rate peaked in the mixed zone near x ≈ 17 m, followed by a rapid decline downstream (Fig. 7). Toward the end of the flume, within the high-velocity zone, the reaction rate showed a slight increase again (Fig. 7).

**Fig 7.**
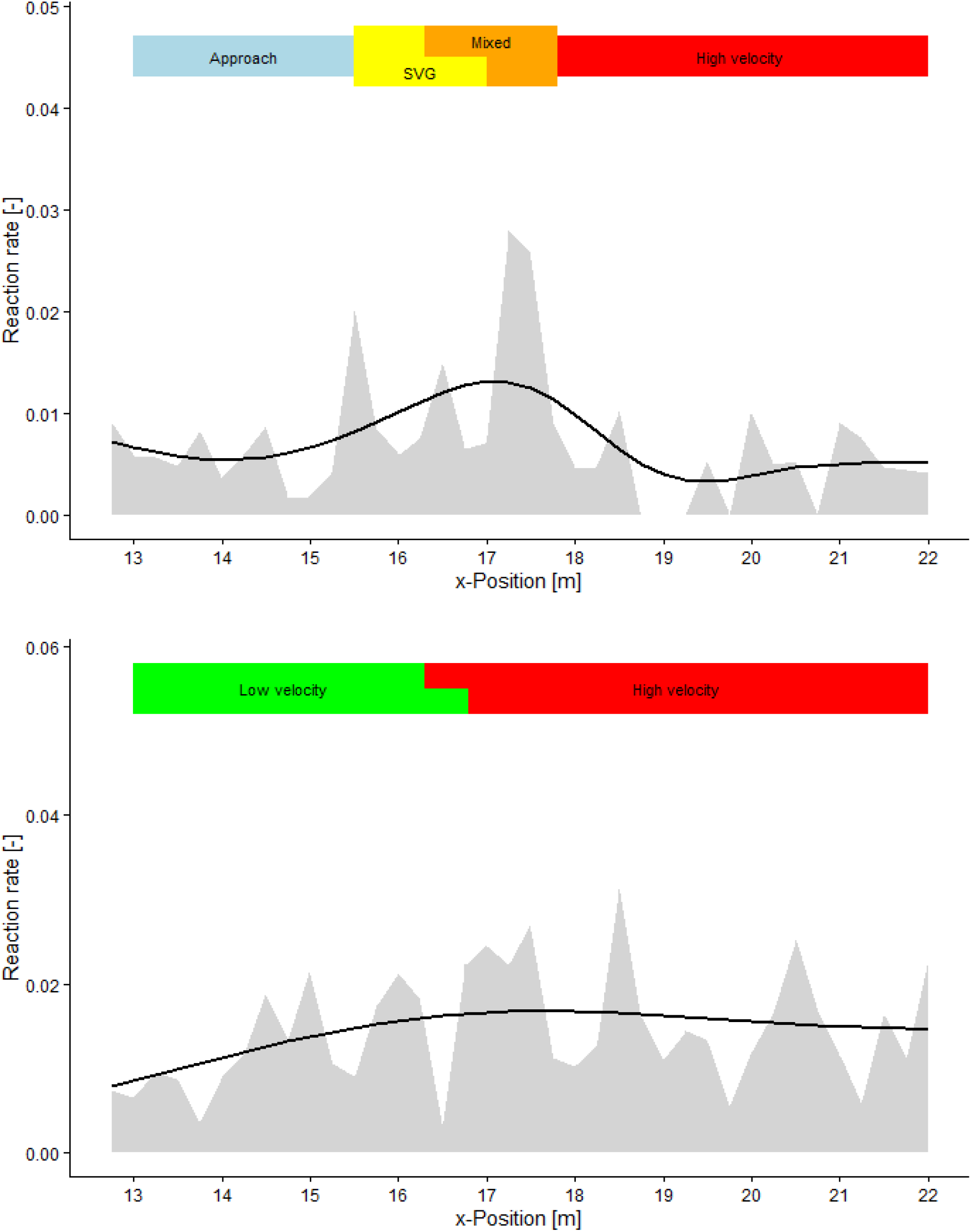
Avoidance reaction rates of fish in daylight (top) and dark (bottom) conditions referred to 25 cm wide flume volume segments (N=38) set virtually along the x-axis of the experimental flume (grey) and fitted GAM function (solid line), the different hydraulic zones along the flume x axis are marked by the colored bar above each graphic, in daylight where the trout specific threshold for SVG of 0,31 m/s/m is available from our experiments the for zones (Approach zone:-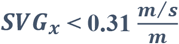; *v*_*x*_ < *u*_*sprint*_; *SVG*_*x*_zone: 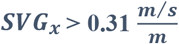; *v*_*x*_ < *u*_*sprint*_; Mixed zone: 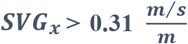; *v*_*x*_ ≥ *u*_*sprint*_; High velocity zone: 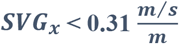; *v*_*x*_ ≥ *u*_*sprint*_) are distinguished, whereas for the dark conditions along the flume x axis only a distinction between low velocity zone *v*_*x*_ ≤ *u*_*sprint*_ and low velocity zone *v*_*x*_ ≥ *u*_*sprint*_ was possible; the end of zones the SVG zone (daylight) or low velocity zone (dark) and the start of the mixed zone (daylight) or high velocity zone (dark) depend on the body length-dependent maximum sprint speed of each fish, this causes a fuzzy boundary between SVG/mixed zone or low/high velocity zone.

In contrast, under dark conditions, reaction rates remained consistently high in the high-velocity zone and decreased downstream only slightly (Fig. 7). Like in the daylight experiments, an increase in reaction rate was observed in the initial flume section, where flow velocities were below the fish’s maximum sprint swimming speed (Fig. 7). Although less pronounced, a reaction rate peak occurred at a similar position as in daylight, coinciding with flow velocities exceeding the fish’s sprint swimming capacity (Fig. 7). Additionally, a notable behavioral difference between light and dark conditions was the significantly higher number of avoidance reactions observed in brown trout in darkness (Fig. 8).

**Fig 8.**
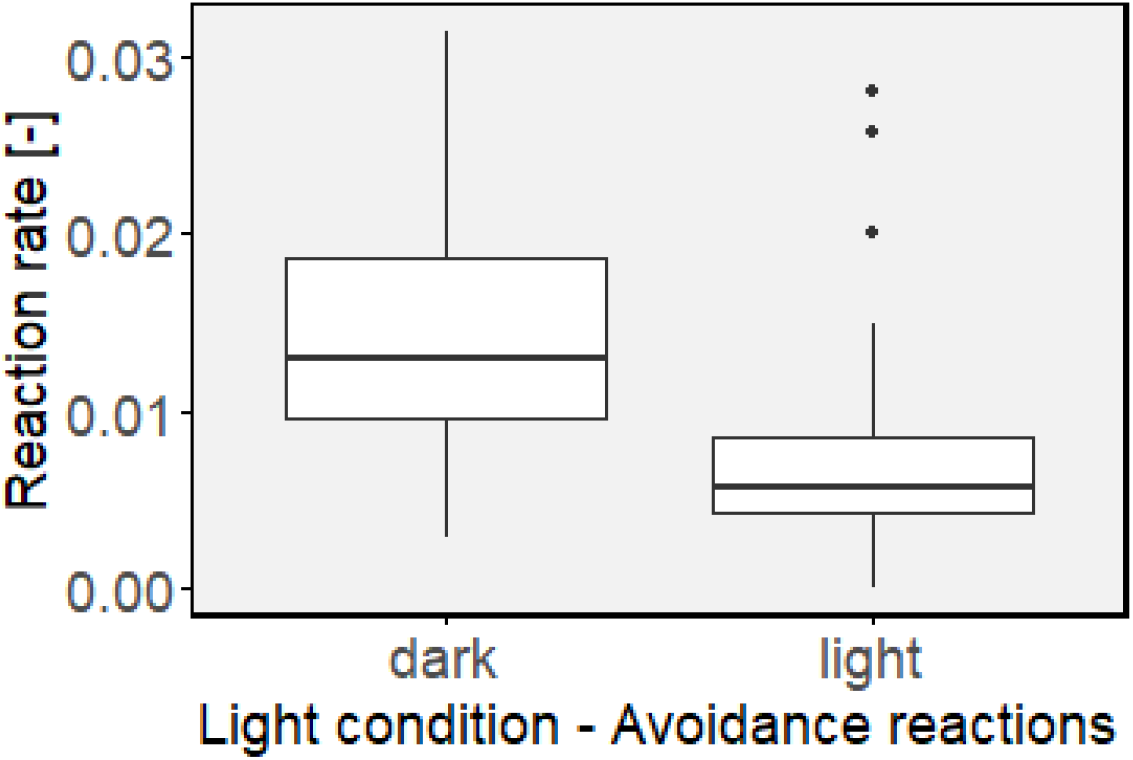
Boxplot showing medians, 25- and 75-percentiles of the avoidance reaction rate within the 25 cm long flume volume segments (N=38) set virtually along the x-axis of the experimental flume (N= 38) in daylight and dark experiments. The data reveal significant differences in reaction rates (Wilcoxon rank sum test) between the fish in dark and daylight avoidance reactions W = 1197, p < 0.001)

Like the spatial pattern of avoidance reactions, the distribution of the six movement modes of fish along the x-axis of the flume differed between daylight and dark experiments (Fig. 10). In daylight, controlled drift, including both strong and weak activity levels, was the predominant movement mode throughout the flume (Fig. 10, Fig. 11). In contrast, under dark conditions, this dominance was less pronounced. Other movement modes, most notably active downstream swimming and drift uncontrolled, were more prevalent across all zones (Fig. 10). In the low-velocity zone, stationary and upstream movements occurred significantly more frequently compared to the daylight trials.

Under daylight conditions fish primarily exhibited controlled drift with weak activity in the approach zone. Already shortly before the start of the SVG zone, the proportion of fish displaying controlled drift with strong activity and even upstream movement increased markedly (Fig 9). In the transition between the mixed and high velocity zone, where flow velocity exceeded the theoretical sprint speed of the test fish (maximum *v*_*sprint*_ =1.94 m/s) and *SVG*_*x*_ surpassing the threshold value for avoidance behavior, many fish switched in the mode *controlled drift* with *strong* swimming activity. (Fig 9). Further downstream a proportion of 41,1 % of the fish first reduced their activity despite the high flow velocity and *SVG*_*x*_ values close to zero, switching in the drift controlled mode from high to weak activity. However further downstream a remarkable proportion of 38,6 % of fish switched again to strong activity. The proportion of the movement modes downstream active and drift uncontrolled was in general very low and negligible in the high velocity zone in daylight conditions (Fig 9).

**Fig 9.**
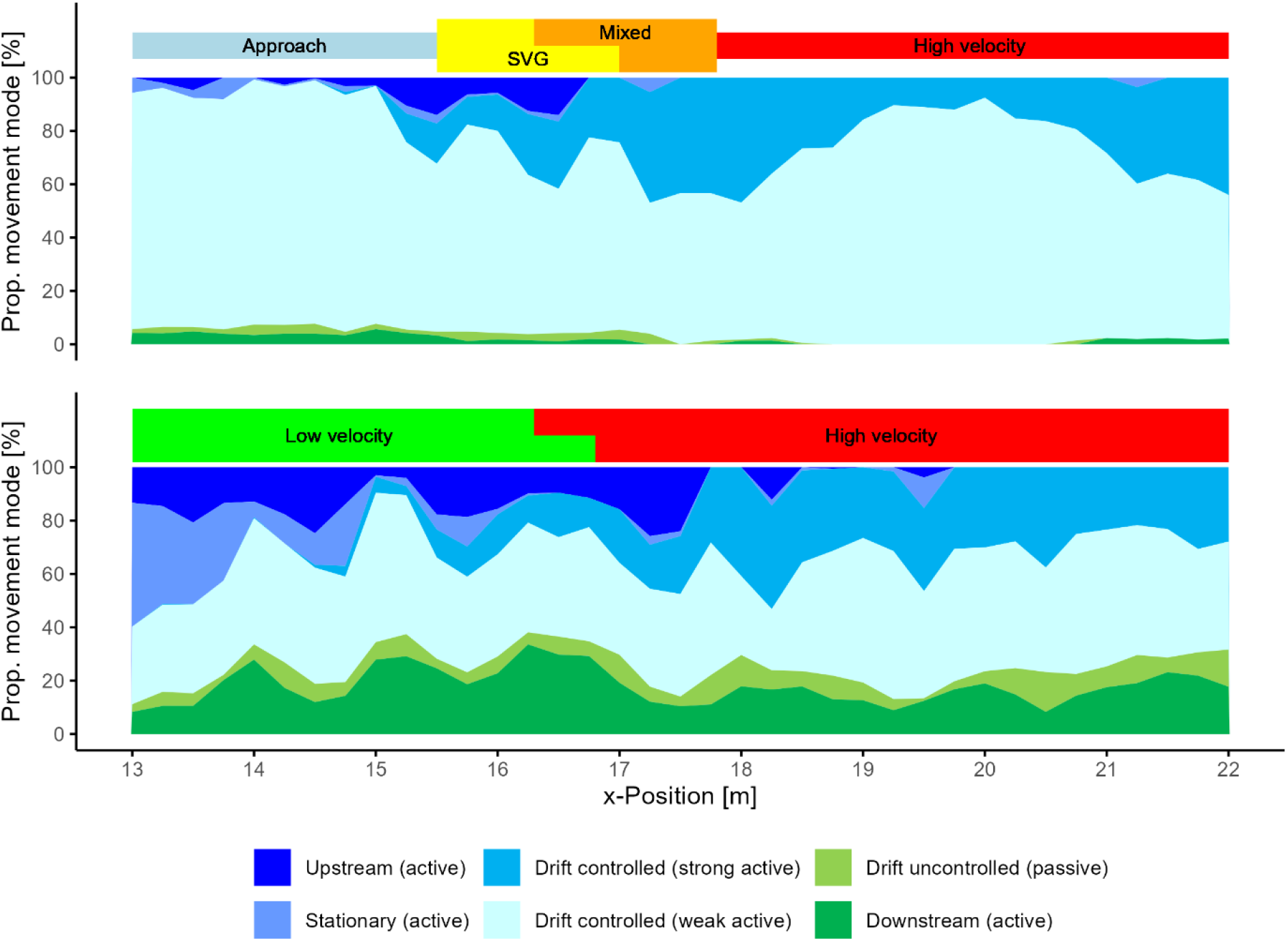
Spatial distribution of the six movement modes along the x-axis of the experimental flume of downstream passing brown trout in daylight (top) and dark conditions (bottom).

Under dark conditions in the low velocity zone, the movement modes active downstream, upstream, and stationary locally accounted for over 15% of the observed movement modes each. However, in the transition to the high-velocity zone the majority shifted into *controlled drift* mode as the high velocity makes upstream movement nearly impossible. A considerable proportion exhibited *strong activity*, indicating an adaptive response to the changing hydrodynamic conditions in dark as well. Remarkably some fish stayed a certain time stationary or even swam upstream in the high flow zone despite the flow velocity exceeding their calculated theoretical maximum. The maximum normed swimming power in dark reached up to 5.7 compared to 3.8 in daylight.

General behavior patterns in the different hydraulic zones and lighting conditions are revealed in movement mode and normed swimming power (Fig 10). The movement mode drift controlled with weak activity with a normed swimming power between 0.04 and 0.085 (Fig 10) was dominant in all zones independently from lighting conditions. However, when the flow velocity exceeded in daylight, the theoretical sprint swimming speed the proportion of fish with strong activity in controlled drift increased to 32.5 % with *P*_*norm*_ of 0.408 in the mixed zone and 26.9 % with *P*_*norm*_ of 0.47 in the high velocity zone. In the dark experiments the proportion of the drift controlled mode with strong activity increased in the high velocity zone to a similar value of 26.8 % but with nearly to the double *P*_*norm*_ of 0.81. The *P*_*norm*_ 0.04 in the drift controlled mode with weak activity was slightly lower than in daylight with 0.052. When flow velocity exceeded the sprint swimming speed of trout, all fish except 1% exhibited controlled downstream drift. Under dark conditions, controlled downstream drift predominated however, active downstream movement (17%), uncontrolled drift (5.2%), and even upstream movement (5.2%) also occurred. Successful upstream movement in the high velocity zone required a mean of *P*_*norm*_ of 3.9 for the fish (Fig 10).

**Fig 10.**
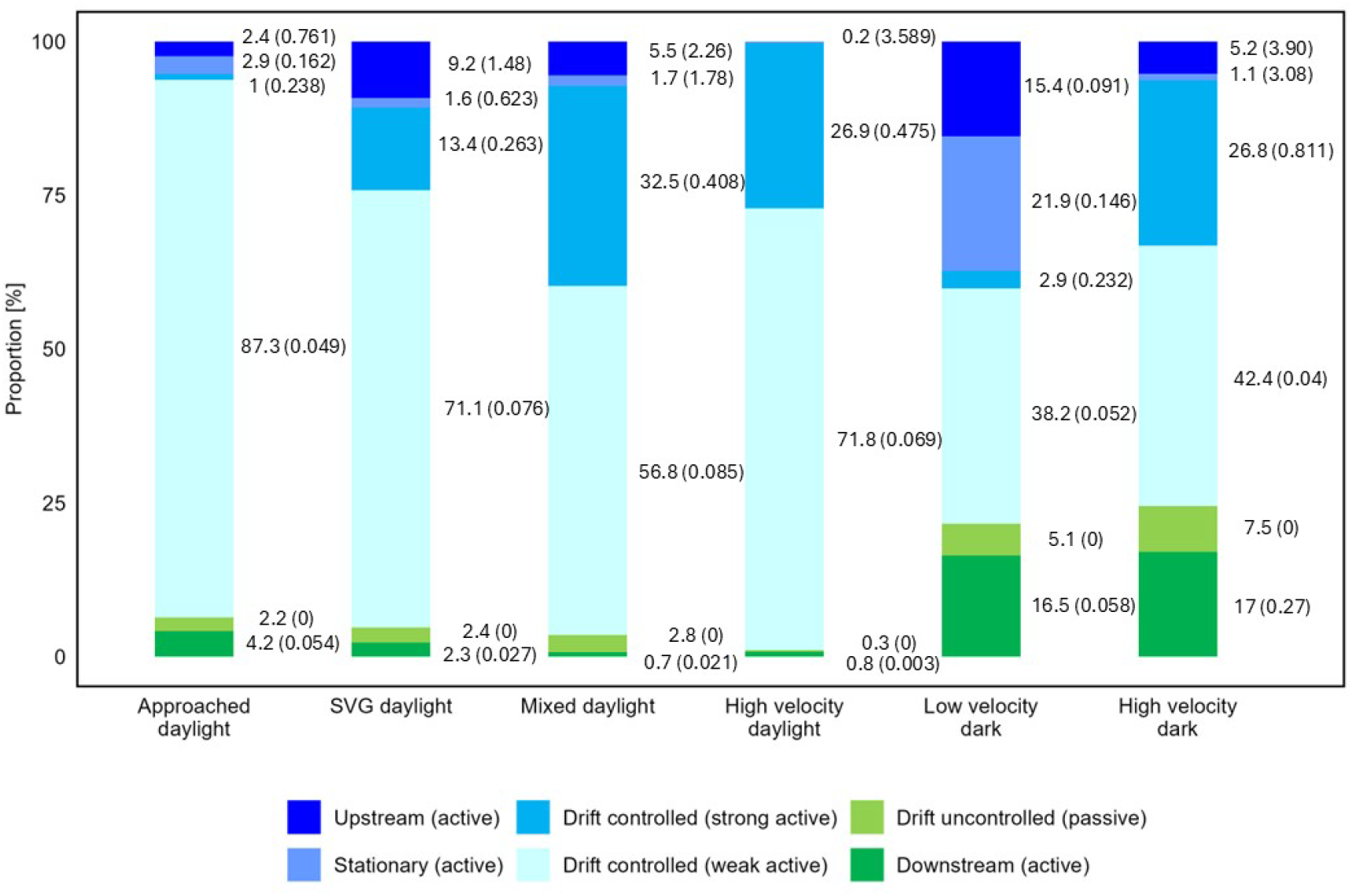
Proportions of the six movement modes in the four hydraulic zones in daylight and in the two in dark conditions along the experimental flume and mean *P*_*norm*_ in brackets of the brown trout in the daylight and dark experiments

## 4. Discussion

This flume study provided insights into the swimming behavior of adult brown trout under hydraulic conditions typical of turbine and pump intakes, particularly in flow regimes with flow velocity exceeding their sprint swimming capacity and high streamwise gradients. By applying a quantitative framework based on five basic movement modes [39,40,61] considering rheotactic orientation, swimming speed and the analysis of swimming power by Hidden Markov Models (HMMs) [52], we were able to detect subtle behavioral changes hard to recognize in common qualitative behavior assessments such as [29,30,34,38,62]. Our findings address critical gaps regarding fish behavior in high-velocity environments [33]. Moreover, the use of measurable variables in behavior analysis allows for high reproducibility of the data and facilitate comparisons across different experimental setups or even fish species. Consequently, this methodology can significantly contribute to a deeper understanding of how fish respond to hydrodynamic conditions and to determine trigger thresholds.

Contradicting results are available in the literature about behavior and rheotactic orientation of downstream migrating Salmonides when approaching hydro power plants (review [63]). Our experiments reflect abstracted but representative flow conditions for hydraulic structures and turbine or pump inlets. These commonly feature high flow gradients, flow accelerations and decelerations, as well as turbulent flow conditions with high flow velocities up to several magnitudes higher than the sprint swimming speed of fish. Under these conditions the majority of brown trout drifted controlled and thus had a positive rheotactic orientation while moving downstream, independently from lighting conditions and streamwise flow velocity below and above the theoretical sprint swimming speed of the fish. When the flow velocity exceeded the trout’s sprint swimming speed, controlled drift was observed in daylight nearly exclusively (> 99 % of observation). This result is in line with those for brown trout [31], rainbow trout [38] and Atlantic salmon kelts [64] with predominantly positive rheotactic oriented fish in accelerated flows. A switch of the orientation to face the flow in connection with active rejection in accelerated flows is a behavior that has already been reported for other salmonids [31,65,66]. Kemp et al. [65] underline that this behavior helps fish to maintain position and control during downstream movement to enable fast avoidance reactions at potentially harmful locations. However, in our dark experiments a proportion of ∼17 % moved negatively oriented actively downstream and ∼7.5 % drifted uncontrolled without a constant rheotactic orientation.

A commonly assumed giving-up behavior models the fish as a passive body in numerical simulations or analytical blade-strike models. Under high velocity conditions, this behavior would imply a switch to negative rheotaxis or a reduction in swimming power in brown trout. However, neither response was observed in our flume experiments, regardless of lighting conditions.

Similar to Pavlov’s findings for juvenile fish [67] the proportion of actively downstream swimming and passive drifting adult trout was higher in the dark compared to daylight. But still the majority of observation showed controlled downstream drifting fish as reported by Haro for Altantic salmon smolts [29] swimming actively against the flow even in the high velocity. In our flume a proportion of fish even exceeded their theoretical maximum swimming power in that condition up to a factor of more than five. Such behavior in a turbine or pump may explain the higher mortality risk of active swimming compared to inactive trout in turbines [23,24]. Although fish can sustain sprint swimming speed for only 10–20 seconds before fatigue [68,69] our results indicate that brown trout approaching the rotor zone of pumps and turbines typically engage in active swimming behavior that reduces their drift speed and thus increases the residence time in the dangerous zone leading to more blade collisions. In consequence this behavioral response, rather than passive drift, increases the likelihood of injury and mortality. Our study results clearly confirm hypothesis (1) that downstream migration fish do not become passive when entering flow zones where the streamwise flow velocities exceed their maximum sprint swimming capacity. Most trout continued to swim actively with a positive rheotactic orientation, even when being subjected to involuntary downstream drift.

Flow velocity and *SVG*_*x*_ were relevant triggers for avoidance behavior reactions of the downstream migrating adult brown trout in our flume experiments. The probability of avoidance reactions by the trout increased significantly at a *SVG*_*x*_ threshold of 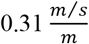 based on the pooled data from all experiments and the daylight data. This threshold value is in-between the mean value described by Vowles & Kemp [31] for avoidance reactions of brown trout in daylight at 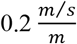 and dark at 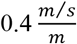. The lower sensitivity to the *SVG*_*x*_ in dark as described by Vowles & Kemp [31] is not supported by our data since the reaction rate of trout in our experiments changed gradually in that condition without a distinct maximum. However, in daylight conditions trout increasingly swam upstream or stayed stationary to avoid further downstream movement when SVG reached 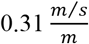. This confirms the initial hypothesis (2) for daylight conditions: Rapid acceleration of the streamwise flow velocity over short spatial distances (high spatial velocity gradients - SVG) induces significant changes in fish swimming behavior.

The observed differences in fish behavior in daylight and dark experiments in response to the SVG indicate that light stimuli function as a secondary signal, enhancing the effectiveness of hydrodynamic cues as proposed by Partan et al. [70]. This interpretation is further supported by an increase in avoidance reaction rates following an initial decline, which occurred only under daylight conditions in the high velocity zone. Given that *SVG*_*x*_ were nearly zero in this flume section, the avoidance reactions here were likely triggered by the high flow velocity causing a fast downstream drift perceivable by the fish visually in daylight. In the dark experiments, the high drift speed may be difficult to detect by the fish due to limited visual cues even when stationing in fish is considered a multisensory process [71]. While fish can perceive SVG through the lateral line organ [72–74] and linear acceleration by the semicircular systems of the inner ear [74], perception of the absolute value of flow velocity is likely more challenging without a visual reference to gauge the ratio of position change relative to their swimming speed [22]. In daylight, fish can track their movement using visual markers [44,75,76], but this sense is likely to be severely restricted in darkness. As a result, a fish in the dark might be less precisely able to detect the speed of downstream drift and adjust swimming speed and swimming direction accordingly. This may explain as well why they more frequently swim actively downstream, drift uncontrolled, or even escape upstream with substantially higher number of avoidance reactions than under daylight conditions.

Regarding hypothesis (3): The pronounced differences in behavioral responses to both SVG and high velocity conditions, together with the markedly higher number of avoidance reactions in darkness, demonstrate that lighting conditions strongly affect how fish respond to spatial velocity gradients and to flow velocities exceeding their maximum sprint swimming capacity. Notably, active downstream swimming and passive drifting were far more common in darkness, even at flow velocities above the sprint threshold, whereas nearly all individuals displayed controlled downstream drifting during daylight. This contrast further confirms the crucial role of lighting conditions in shaping fish movement behavior under challenging hydraulic situations.

The divergent behavior patterns found for brown trout in daylight and dark are relevant for the integration of behavior rules into models. The fact that the majority of brown trout orientated positive rheotactic aligned with the main flow vector independently from SVG, flow velocity and lighting conditions can be implemented in blade strike models [22,77]. For models where the swimming activity of the fish can be implemented [78] the following results may be relevant. In daylight the movement mode drifted controlled caused a delay of the downstream movement compared to the flow velocity dominated through the flume, independently from SVG or flow velocity. When the flow velocity exceeded the theoretical sprint swimming speed of fish the controlled drift mode showed a mean swimming speed in daylight of above 40% of theoretical sprint swimming speed (*P*_*norm*_ 0.4) and even slightly above 80 % (*P*_*norm*_ 0.8).

Given that adult brown trout are primarily nocturnal [79], and that closed conduits in hydraulic facilities are rather dark environments, the behavioral patterns observed under dark conditions may be more relevant than those carried out in illuminated conditions. In any case, the lighting condition should be considered in behavioral models for corresponding conditions.

The transfer of these ethohydraulic laboratory findings into real-world applications will require further validation in field studies to assess their practical applicability and reliability. Given the lack of detailed behavioral data for various fish species, establishing effective downstream passage systems at hydropower plants remains a challenging task [28]. To address this, ongoing experiments with additional species will expand the behavioral rules established here to include representatives of Cyprinoidea and Percidae, further enhancing the applicability of our findings.

## 5. Conclusion

Based on the quantitative analysis of movement modes and swimming activity using HMM approaches in this study, we gained detailed fish response data and valuable insights into fish behavior under challenging flow conditions with flow velocities exceeding the sprint swimming speed of brown trout. Our results demonstrate the influence of spatial velocity gradients (*SVG*_*x*_) and flow velocities on the behavior of adult brown trout (*Salmo trutta*) during downstream migration in hydraulic environments similar to turbine and pump intakes. The experiments revealed that *SVG*_*x*_ serves as a trigger for changes in swimming behavior, independent of the lighting condition, with a threshold of 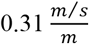 for an increase of avoidance reactions in daylight. However, lighting conditions significantly influenced the frequency and nature of fish responses, with more reactions and greater behavioral variability in darkness generally in all hydraulic conditions. Fish preferred distinct movement modes depending on hydraulic conditions, with increased controlled drifting and higher swimming power as flow velocities approached or exceeded their sprint swimming capacity. They did not give up. The majority of brown trout remained rheotactically positively oriented and delayed downstream drift when the flow velocity surpassed their sprint swimming speed, independently from lighting condition. These findings underscore the importance of considering fish behavior and the modulating effect of environmental factors such as lighting condition when modeling mortality risks and designing fish-friendly hydraulic infrastructure. Integrating behavioral rules comprising swimming mode and swimming power derived from this study into predictive risk assessment models could enhance the development of methods for safer hydropower systems and pumping stations, reducing the negative ecological impact of human alterations in the water bodies. Future research should validate these findings at real world hydraulic machinery sites and expand the scope to include other fish species.

## Acknowledgments

The study was part of the RETERO project funded by the German Federal Ministry of Research and Education under the program “Alternative Methods to Animal Experiments” with the funding references 031L0152B, 16LW0168, 031L0152A, D 161L0152B.

The authors like to thank a group of people who were deeply involved in the project: N. Mueller (IWD) for her work in the first phase of the project; D. Powalla (OVGU) for his support during the flume design phase. L. Backhaus (IWD) for the software to automatically cut the high number of raw-data-videos; M. Geyer, O. Just, T. Graefe and M. Ehrig (all laboratory staff, IWD) for the construction and operation of the laboratory model, as well as all student assistants (IWD, IGF) which helped everywhere a hand was needed. Special thanks to the Thuringian fisheries administration (TMIL) for their support and Marcus Wendler for “renting” the brown trout for the experimental period. We would like to sincerely thank the anonymous reviewer for their constructive and insightful comments on the first version of this manuscript.

## Author Contributions

**Conceptualization:** Falko Wagner, Ianina Kopecki, Stefan Hoerner, Tom Roessger

**Data curation:** Falko Wagner, Ianina Kopecki, Kathrin Maltzahn, Tom Roessger

**Formal analysis:** Falko Wagner, Ianina Kopecki, Jelger Elings, Mansour Royan, Márcio S. Roth, Tom Roessger, Ulf Enders

**Funding acquisition:** Falko Wagner, Jürgen Stamm, Stefan Hoerner

**Investigation:** Falko Wagner, Andreas Lindig, Kathrin Maltzahn, Tom Roessger

**Methodology:** Falko Wagner, Ianina Kopecki, Jelger Elings, Márcio S. Roth, Stefan Hoerner, Tom Roessger

**Project administration:** Falko Wagner, Jürgen Stamm, Stefan Hoerner, Tom Roessger

**Resources:** Falko Wagner, Ianina Kopecki, Jürgen Stamm

**Supervision:** Falko Wagner, Jürgen Stamm

**Validation:** Falko Wagner, Ianina Kopecki, Tom Roessger

**Visualization:** Falko Wagner, Ianina Kopecki, Mansour Royan, Márcio S. Roth, Tom Roessger, Ulf Enders

**Writing – original draft preparation:** Falko Wagner

**Writing – review and editing:** Falko Wagner, Ianina Kopecki, Jelger Elings, Márcio S. Roth, Stefan Hoerner, Tom Roessger

